# BindCORE: Biophysical Ensemble Learning for Predicting Interaction Sites in Intrinsically Disordered Regions

**DOI:** 10.64898/2026.08.20.746047

**Authors:** Nicolas Buton, Luiz Felipe Piochi, Hamed Khakzad

## Abstract

Intrinsically disordered proteins and regions (IDPs/IDRs) mediate diverse cellular functions through binding segments whose functional properties are encoded in dynamic conformational ensembles rather than a single static state. Existing predictors of linear interacting peptides (LIPs) and molecular recognition features (MoRFs) rely primarily on sequence-derived features, leaving ensemble-level biophysical properties largely unexplored. Here, we introduce BindCORE, an ensemble-aware deep learning framework that integrates global, local, and pairwise biophysical descriptors to predict interaction sites within IDRs. These features are processed through a multi-scale architecture that enables information exchange between sequence- and ensemble-based global, local, and pairwise information. Across established LIP and MoRF benchmarks, BindCORE consistently improves performance over sequence-based baselines, demonstrating the predictive signals of ensemble-derived properties beyond sequence-based representations alone. Feature-attribution analyses reveal that pairwise descriptors are the dominant contributors to prediction, while solvent accessibility, backbone dihedral entropy, and global geometric properties provide complementary information. Feature-importance rankings vary substantially across ensemble flavours, indicating that different conformational generators encode distinct biophysical signatures of interaction-site propensity. Together, our results show that conformational ensembles contain interpretable determinants of LIP and MoRF binding residues and establish BindCORE as a general framework for incorporating biophysical information into the prediction of functional regions in intrinsically disordered proteins. BindCORE is freely available as a ready-to-use Google Colab notebook (BindCORE Colab notebook).

**Key Messages:**

- BindCORE integrates ensemble-derived biophysical descriptors to predict residue-level interaction sites in intrinsically disordered proteins.
- Ensemble-derived features improve prediction performance over state-of-the-art sequence-based methods on both LIP and MoRF benchmarks.
- Pairwise ensemble descriptors, especially contact and dynamic cross-correlation maps, provide the strongest signals for predicting interaction-site residues, while global chain geometry, solvent accessibility, and backbone dihedral preferences add complementary information.

## 1 Introduction

Intrinsically disordered proteins and regions (IDPs/IDRs) challenge the traditional sequence-structure-function paradigm by existing as highly dynamic, structurally heterogeneous conformational ensembles under physiological conditions [1–3]. Their biological functions, ranging from cellular signaling to regulatory network modulation, are inextricably linked to this conformational plasticity [4–6]. A key mechanism by which IDPs and IDRs exert their functions is through linear interacting peptides (LIPs), a broad class of binding segments that includes Molecular Recognition Features (MoRFs), short linear motifs (SLiMs), and protean segments (ProSs). While SLiMs are typically very short (∼3–10 residues), MoRFs and ProSs generally span longer segments (∼10–25 residues or more). Embedded within disordered regions, these elements enable diverse molecular interactions. MoRFs and ProSs primarily mediate protein–protein interactions and often undergo a disorder-to-order transition upon binding, whereas SLiMs exhibit broader interaction profiles, engaging not only proteins but also nucleic acids, lipids, and small molecules. More generally, LIPs and ProSs encompass a range of binding behaviors depending on their precise definitions, and many undergo coupled folding and binding upon interaction with their physiological partners [7–9]. LIPs, MoRFs, and related binding residues mediate many protein–protein interactions underlying human disease, making their accurate identification a problem of both fundamental and translational importance [10]. Because the binding behavior of these regions is dynamically encoded within the broader sampled phase space of the protein rather than in a single native state, understanding and predicting these functional motifs fundamentally requires an ensemble-centric, sequence-ensemble-function perspective [11–13].

Despite the inherently dynamic nature of IDPs, existing computational methods for predicting LIPs, MoRFs, and related binding residues rely predominantly on sequence-derived features. Tools such as ANCHOR [14] and MoRFpred [15] exploit evolutionary profiles, physicochemical propensities, and predicted disorder scores. More recent models, including CLIP [16] and MoRFchibi 2.0 [17], extend this paradigm through deeper architectures and more extensive feature engineering, yet remain fundamentally sequence-based. In contrast, structure-informed approaches such as FLIPPER [18] incorporate three-dimensional information to identify interaction-prone regions; however, they rely on experimentally determined structures of the full complex, including the binding partner. This requirement severely restricts their applicability and makes it impractical to explore the vast space of potential interactions. Consequently, no existing method systematically leverages ensemble-level biophysical descriptors derived from the IDP conformational landscape, leaving underexploited the structural heterogeneity that governs interaction propensity in solution and motivating the development of approaches that do not depend on prior knowledge of binding partners. Historically, this omission was driven by the prohibitive cost of obtaining high-quality structural ensembles. The field is now experiencing a rapid expansion in ensemble generation capabilities [19]. On the physics-based side, advanced all-atom force fields [20, 21] and all-atom methods such as bAIes [22] can now sample IDP conformations at scale. Specialized coarse-grained models, including CALVADOS [23, 24], Mpipi-GG [25], and Martini3-IDP [26], also enable large-scale IDP sampling. On the machine-learning side, a new generation of generative models predict conformational ensembles directly, including IDP-specialized tools such as IDPFold2 [27], IDPForge [28], and STARLING [29], as well as flow-matching approaches such as AlphaFlow and ESMFlow [30].

This wealth of structural data creates a new computational bottleneck. Feeding raw unaligned ensembles of varying sizes directly into deep learning models is an immensely challenging task, heavily prone to noise and often prohibitive given limited computational resources [31]. To bridge this gap and to fully harness structural dynamics for functional prediction, we propose a two-tiered computational framework. First, we introduce EnsembleMDP, an open-source library that systematically extracts biophysically meaningful descriptors from conformational ensembles. Instead of relying on raw atomic coordinates, EnsembleMDP summarizes ensemble behavior through a diverse set of structural and dynamical properties spanning global, residue-level, and pairwise features. Importantly, the framework captures not only average conformational behavior but also ensemble heterogeneity through variation and entropy-based statistics, enabling a compact yet information-rich representation of protein dynamics [32, 33]. Building on these representations, we introduce BindCORE, an ensemble-aware deep learning architecture that leverages ensemble-derived biophysical features to predict binding residues, including LIP and MoRF regions, at single-residue resolution. To our knowledge, BindCORE is the first deep learning framework to explicitly integrate conformational ensemble information for residue-level binding-region prediction in IDPs.

## 2 Materials and Methods

To evaluate whether conformational ensembles encode information relevant to binding-region prediction beyond sequence alone, we developed a workflow that transforms ensembles generated by diverse computational approaches into quantitative biophysical descriptors using EnsembleMDP. These descriptors were then incorporated into BindCORE for residue-level prediction of LIP and MoRF binding regions. An overview of the pipeline is shown in Figure 1a, and the individual steps are described below.

**Figure 1.**
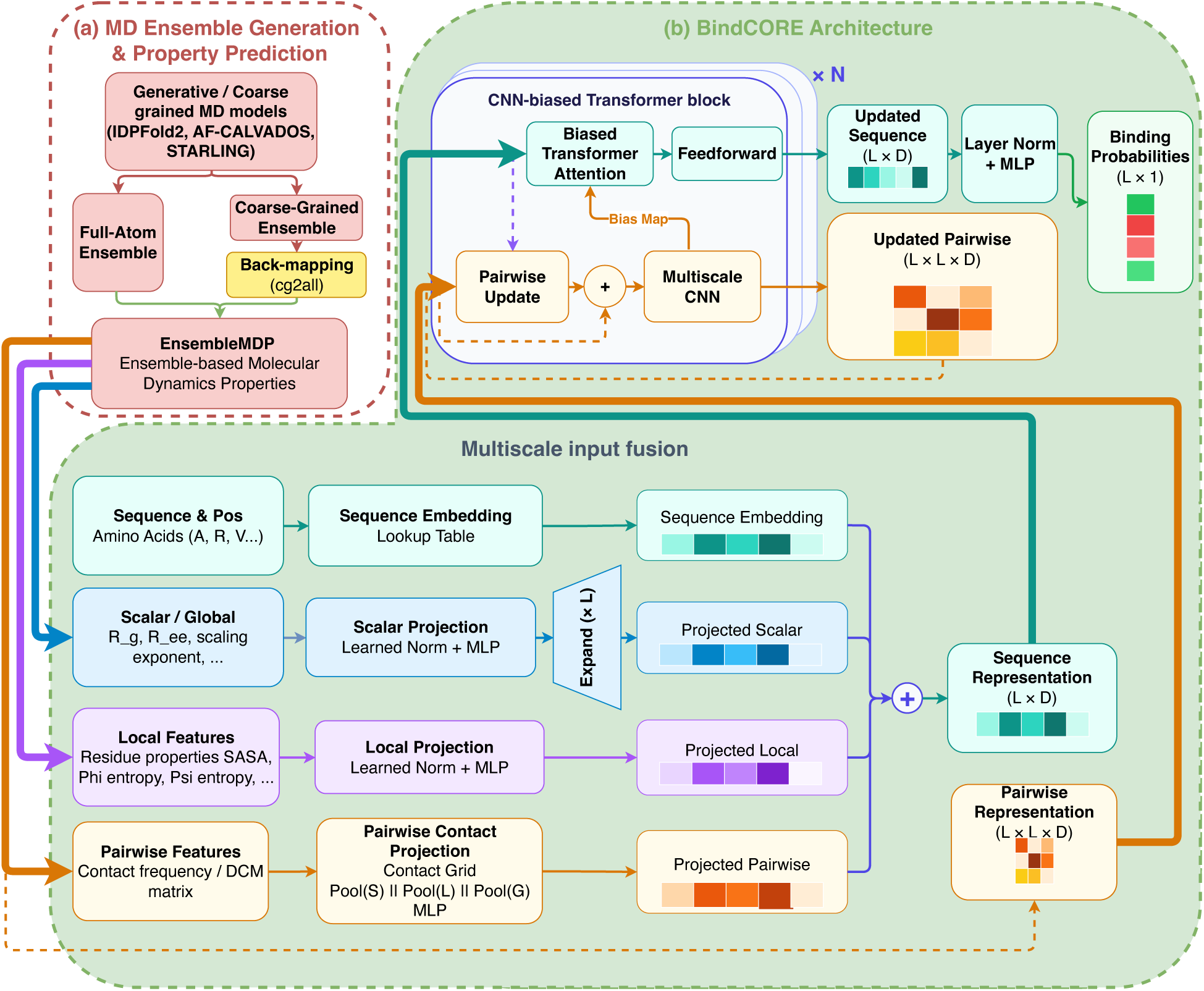
Integrated EnsembleMDP and BindCORE workflow. **(a)** Ensemble-processing workflow: conformational ensembles are generated using physics-based, hybrid, or machine-learning approaches; coarsegrained conformations are backmapped to full-atom structures with cg2all using center-of-mass inputs for AF-CALVADOS and *C_α_*-trace inputs for IDPFold2 and STARLING; and EnsembleMDP converts the resulting ensembles into interpretable global, residue-level, and pairwise descriptors. **(b)** BindCORE integrates sequence, global, local, and pairwise ensemble features into residue embeddings; CNN-biased Transformer blocks jointly update residue and pairwise representations, and a final MLP outputs residue-level binding probabilities.

### 2.1 Conformational generation and geometric validation

To generate conformational ensembles, we selected IDPFold2, AF-CALVADOS, and STARLING, given their strong agreement with experimental validation data, particularly SAXS- and NMR-derived observables, across different benchmarks of protein ensemble prediction [22, 27, 29, 34], while remaining computationally tractable for application to thousands of proteins. For each tool, we used the default number of generated conformations: 100 frames for IDPFold2, 400 frames for STARLING, and 1,000 frames for AF-CALVADOS. As all three methods generate coarse-grained conformational ensembles, every frame was backmapped to full-atom coordinates using cg2all [35] before giving as input to EnsembleMDB (Figure 1a). AF-CALVADOS ensembles were backmapped with the cg2all model trained for center-of-mass coarse-grained inputs, whereas IDPFold2 and STARLING ensembles were backmapped with the cg2all model trained for *C_α_*-trace inputs. Up to 1,000 randomly selected frames per protein were quality-checked when available. This included covalent bond lengths, backbone N–*C_α_*–C angles, steric clashes after excluding bonded neighbors, peptide-bond planarity through *ω* deviations, and Ramachandran outliers. Frames were considered geometrically viable only if they passed all criteria, using a maximum 1% bad-bond ratio, a 0.7 van der Waals overlap factor for clashes, and a maximum 5% bad *ω* or Ramachandran residues. These diagnostics were used to compare reconstruction quality across ensemble sources and to guide interpretation of downstream results, but no prediction benchmark was restricted to only fully viable frames.

### 2.2 EnsembleMDP framework

To characterize the conformational landscape of each protein ensemble, we calculated biophysical descriptors grouped into global, residue-level (local), and pairwise features, completing the property-extraction step shown in Figure 1a. For frame-dependent properties, summary statistics across the *N* conformations were retained, including the mean and standard deviation and, where applicable, entropy-based measures. This representation converts heterogeneous coordinate ensembles into fixed-format inputs suitable for residue-level prediction while preserving interpretable biophysical information.

#### 2.2.1 Global structural descriptors

Global descriptors summarize the polymer-scale behavior of each conformation and are particularly informative for IDPs, whose biological activity often depends on shifts between expanded, collapsed, and anisotropic states. Chain extension was quantified using the end-to-end distance (*R_ee_*), defined as the Euclidean distance between the *C_α_* atoms of the first (**r**_1_) and last (**r***_L_*) residues:

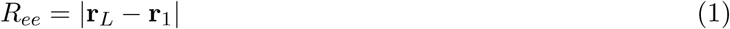

We retained both the mean and standard deviation of *R_ee_*^2^ across the ensemble, together with the mean and standard deviation of the maximum intrachain *C_α_*–*C_α_* diameter, to capture chain extension independently of terminal-residue placement.

Overall compaction was measured using the radius of gyration (*R_g_*), which describes how atomic mass is distributed around the center of mass (**r***_cm_*):

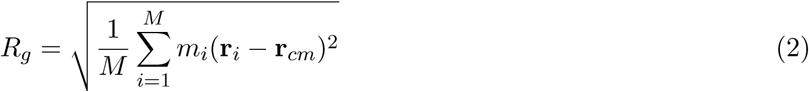

where *m_i_* is the mass of atom *i* and *M* is the total mass. In contrast to *R_ee_*, which depends only on the terminal residues, *R_g_* captures the full chain and is therefore less sensitive to fluctuations or atypical positions at the end terminals.

To complement chain size, we characterized ensemble shape using the gyration tensor **S**:

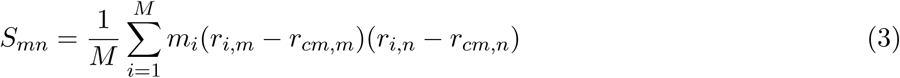

where *m, n* ∈ {*x, y, z*}. Diagonalization of **S** yields eigenvalues *λ*_1_ ≥ *λ*_2_ ≥ *λ*_3_, from which we derived descriptors that distinguish spherical, elongated, flattened, and anisotropic conformations:

- **Principal gyration eigenvalues:** the mean and standard deviation of *λ*_1_, *λ*_2_, and *λ*_3_, which quantify ensemble size along the major, intermediate, and minor principal axes.
- **Principal-axis ratios:** the mean and standard deviation of *λ*_1_*/λ*_2_, *λ*_1_*/λ*_3_, and *λ*_2_*/λ*_3_, which capture elongation and flattening independently of absolute scale.
- **Asphericity (***b*): deviation from a spherical mass distribution, 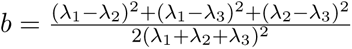.
- **Prolateness (***S*): discrimination between elongated (*S >* 0) and oblate (*S <* 0) conformations.
- **Normalized Acylindricity (***c_norm_*): asymmetry between the two minor principal axes, *c_norm_* = 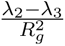.
- **Relative Shape Anisotropy (***κ*^2^): scale-free summary of the overall departure from isotropy, *κ*^2^ = 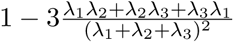.
- Finally, we included the Flory scaling exponent *ν*to relate global dimensions to polymer-physics regimes. Under the power-law relationship *R_g_* ≈ *R*_0_*L^ν^*, where *L* is the sequence length, and *ν* distinguishes compact from expanded-chain behavior. We estimated *ν* through linear regression in log-space, ln(*R_g_*) = *ν* ln(*L*) + ln(*R*_0_), where ln(*R*_0_) is the intercept.

#### 2.2.2 Residue-level descriptors

Residue-level descriptors localize the analysis along the sequence and are particularly relevant for binding-residue prediction because interaction-prone regions often differ from the surrounding disordered background in flexibility, pre-structuring, and solvent exposure. Local conformational heterogeneity was quantified using the Shannon entropy of the backbone torsion angles *ϕ* and *ψ*, computed independently for each residue. Each angle was histogrammed over [−*π, π*] using 60 equal-width bins of 6^◦^, and the marginal entropy was computed as:

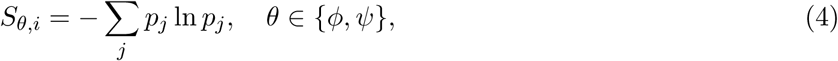

where *p_j_* is the fraction of valid frames falling in bin *j* for a given residue and torsion angle. This yields two independent per-residue entropy tracks. Higher values indicate broader marginal sampling of a given torsion angle, whereas lower values indicate torsional restriction and possible transient pre-organization. Because *ϕ* and *ψ* are treated independently, correlations between the two angles—as would be captured by a joint Ramachandran entropy—are not reflected in these quantities.

Solvent exposure was represented by solvent accessible surface area (SASA), computed with a 1.4 Å probe. For each residue, we retained the ensemble mean and standard deviation of both absolute SASA and relative SASA, to distinguish constitutively exposed residues from positions undergoing repeated burial–exposure transitions while accounting for residue-type-specific maximum accessibility.

Transient pre-structuring was captured by converting DSSP assignments into eight-state propensities for *β*-bridge (B), extended strand (E), 3_10_ helix (G), *α* helix (H), *π* helix (I), coil (C), bend (S), and turn (T), yielding a residue-wise profile of secondary-structure preferences across the ensemble.

#### 2.2.3 Pairwise descriptors

Pairwise descriptors complement residue-wise measurements by capturing transient intramolecular organization and long-range coupling across the ensemble. We considered three pairwise quantities: dynamic cross-correlation, distance fluctuation, and contact frequency.

Correlated residue motion was quantified using the dynamic cross-correlation matrix (DCCM), computed from the *C_α_* displacement vectors across the ensemble:

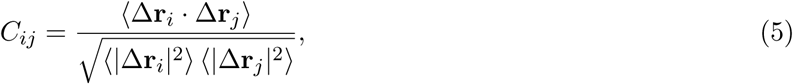

where Δ**r***_i_* = **r***_i_* −⟨**r***_i_*⟩ denotes the displacement vector of residue *i* relative to its ensemble-averaged position. Positive values of *C_ij_* indicate concerted motion between residue pairs, whereas negative values indicate anti-correlated motion.

Distance fluctuations, also referred to as communication propensity [36], were defined as the standard deviation of the distance between the *C_α_*atoms of residues *i* and *j*:

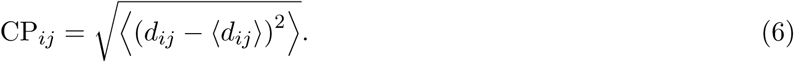

Here, *d_ij_* is the *C_α_*–*C_α_* distance between residues *i* and *j*. Low CP*_ij_* values indicate that a residue pair maintains a relatively stable separation across the ensemble, consistent with stronger mechanical coupling and potential allosteric communication.

We additionally computed the contact frequency map, defined as the probability *P_ij_* that residues *i* and *j* remain within an 8 Å cutoff across the ensemble. This descriptor captures recurrent transient contacts that contribute to conformational compaction and the formation of binding-competent subensembles.

### 2.3 BindCORE architecture

BindCORE is a multi-scale Transformer that jointly processes four complementary protein feature modalities to produce per-residue binding-site predictions. The model was designed to combine sequence identity with ensemble-derived structural information at three resolutions: protein-level descriptors of global conformational state, residue-level descriptors of local environment and flexibility, and pairwise descriptors of transient intramolecular organization. Figure 1(b) provides an overview of the architecture, whose main components are detailed below.

#### Multi-scale input fusion

Given a protein of length *L*, four feature streams are projected independently into a shared embedding space ℝ*^E^* and summed to form the initial residue representation *H*^(0)^ ∈ ℝ*^L^*^×^*^E^*. Amino-acid sequence tokens are encoded with a learned embedding and optional sinusoidal positional encoding. Per-residue local features, including secondary-structure propensities, backbone torsional descriptors, and solvent-accessibility features, are normalized and projected to ℝ*^E^* with a two-layer MLP. Per-protein scalar features, including chain-size and shape descriptors such as *R_g_*, *R_ee_*, gyration-tensor summaries, and the scaling exponent, are normalized, projected to ℝ*^E^*, and broadcast across the *L* residue positions. Pair-wise features *P* ∈ ℝ*^C^*^×^*^L^*^×^*^L^*, including contact-frequency, distance-fluctuation, and dynamic cross-correlation maps, are compressed into per-residue context vectors using three-scale row pooling: short-range averages over neighboring residues, long-range averages outside the local window, and global row averages over the full sequence. The concatenated pooled pairwise summaries are then projected to ℝ*^E^*, giving each residue an initial representation of its contact and coupling environment.

The fused input is defined as

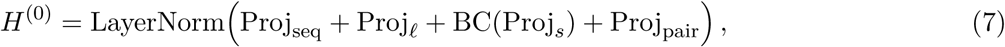

where BC(·) denotes broadcasting over sequence positions. Layer normalization after summation places the heterogeneous streams on a common scale before sequence processing.

#### CNN-biased Transformer blocks

The fused representation is processed by *N* CNN-biased Transformer blocks. Each block maintains and refines two coupled representations: a residue embedding *H* and a pairwise map *P* . This design allows sequence-level evidence and ensemble-derived pairwise organization to exchange information throughout the network rather than being combined only at the input layer.

First, the current residue representation updates the pairwise stream through a low-dimensional outer-product operation inspired by Evoformer-style coupling [37]. Residue embeddings are projected into a compact latent space, pair-specific interactions are formed by an outer product, and the resulting update is symmetrized, normalized, and added residually to *P* . A depthwise-separable 3 × 3 convolution further refines the pairwise map, allowing local spatial patterns in the contact or correlation representation to be smoothed and propagated.

Second, a multi-scale pairwise CNN converts the refined pairwise map into an attention-bias tensor. The encoder applies channel-specific spatial filtering followed by parallel dilated convolutional branches with dilation rates selected during hyperparameter tuning, capturing both local contacts and broader long-range structural dependencies. Branch outputs are merged, normalized, passed through a GELU nonlinearity, and projected to a per-head bias 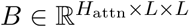 for multi-head attention.

Third, standard scaled dot-product attention [38] over residue embeddings is augmented with this pair-wise structural bias:

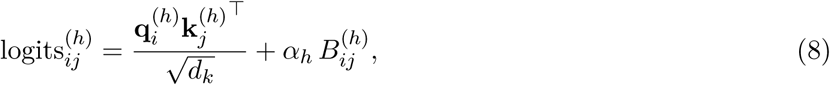

where *α_h_* is a learnable per-head gate initialized to zero. This makes the block equivalent to ordinary self-attention at the beginning of training, allowing structural biases to be introduced gradually as the model learns when pairwise ensemble information is useful. Padding positions are masked in both query and key directions. A position-wise feed-forward network with pre-normalization, GELU activation, dropout, and residual connections completes each block. This bidirectional coupling allows contacts, correlated motions, and distance fluctuations to continuously shape residue-level representations, while the residue representation, in turn, refines the pairwise stream. BindCORE therefore combines Evoformer-like iterative sequence– pair interaction with explicit multi-scale convolutional processing of ensemble-derived pairwise maps.

#### Residue-level classification

After the final block, each residue embedding *H_i_*^(^*^N^*^)^ ∈ ℝ*^E^* is passed through a LayerNorm and an MLP classification head consisting of a linear layer, GELU activation, dropout, and final linear projection to produce a residue-level binding logit. Sequence masks exclude padded positions from the loss. Separate models were trained end-to-end with binary cross-entropy for the LIP and MoRF prediction tasks.

### 2.4 Selected hyperparameters

Hyperparameters for the six BindCORE variants (LIP and MoRF prediction using IDPFold2, AF-CALVADOS, or STARLING ensembles) were optimized independently with a Tree-structured Parzen Estimator (TPE) over 100 trials (Supplementary Figure S1). Despite independent optimization, a clear pattern emerged across ensemble sources. Models trained on IDPFold2 and AF-CALVADOS consistently selected rich feature sets combining local residue descriptors (e.g., backbone geometry, solvent accessibility, and secondary-structure propensities), pairwise interaction features, and sequence embeddings. In contrast, both STARLING-based models converged to compact scalar-only representations augmented with positional embeddings, suggesting that global ensemble statistics provide more reliable information than local structural descriptors for these ensembles. Architecturally, all variants favored shallow networks with only 3–4 transformer blocks, indicating that performance differences primarily arise from the learned representations rather than increased model capacity. The strongest distinction was in regularization: IDPFold2- and AF-CALVADOS-based models consistently required only modest regularization, whereas STARLING-based models favored substantially stronger regularization, reflecting a more constrained optimization landscape.

#### 2.4.1 BindCORE Studio

To facilitate application to user-provided sequences, we developed BindCORE Studio (Supplementary Figure S2), a cloud-based interface available through Google Colab, with an equivalent local Jupyter notebook implementation. Users provide one or more protein sequences in FASTA format, and the pipeline automatically performs (i) conformational ensemble generation using IDPFold2, (ii) backmapping to all-atom structures with cg2all, (iii) ensemble feature extraction using EnsembleMDP, and (iv) binding-region prediction with BindCORE for both LIPs and MoRFs. The interface returns per-residue binding propensity profiles together with summary statistics describing the conformational ensemble, including the predicted binding regions, the mean radius of gyration (*R_g_*), and the Flory scaling exponent (*ν*).

### 2.5 Benchmark datasets and evaluation

BindCORE was evaluated on two residue-level binding-site prediction tasks: the LIP benchmark from CLIP [16] and the MoRF benchmark from MoRFchibi 2.0 [17]. For the primary evaluation, proteins longer than 1024 residues were excluded to keep conformational-ensemble generation computationally tractable. A second evaluation on proteins shorter than 380 residues enabled comparison with STARLING-based models, whose current inference is limited to this sequence length. Sequence-based baselines were evaluated on the same benchmark subsets. After filtering, the MoRF benchmark comprised 608 training proteins and 125 test proteins, whereas the LIP benchmark contained 806 training proteins and 332 test proteins. The two datasets differ substantially in their annotation characteristics: MoRF regions are generally longer and more abundant, whereas LIP regions are short and constitute a highly imbalanced positive class. A detailed statistical summary of both datasets is provided in Supplementary Table S1. To prevent performance inflation due to sequence leakage, both benchmarks were originally constructed using strict homology-aware train–test partitioning. The MoRF benchmark enforces less than 30% sequence identity between training and test proteins and further masks locally homologous regions using Homology Alignment Masking (HAM) [17]. The LIP benchmark was partitioned using CD-HIT with a 25% sequence identity threshold [16]. Performance was evaluated at the residue level using Matthews correlation coefficient (MCC), F1 score, precision, recall, area under the receiver operating characteristic curve (AUC-ROC), and average precision (AP). Because both tasks involve highly imbalanced positive classes, AP was used as the primary ranking metric, while AUC-ROC is also reported to facilitate comparison with previous studies.

### 2.6 Residue-level analysis of ensemble-derived properties

To determine whether EnsembleMDP descriptors distinguish annotated binding residues before model training, we compared descriptor distributions between binding and non-binding residues. Violin plots were generated for each dataset–ensemble-source combination: LIP and MoRF annotations paired with IDPFold2, AF-CALVADOS, and STARLING ensembles. Local properties were analyzed directly because EnsembleMDP reports them at residue resolution. Pairwise properties were first converted from *L* × *L* maps to residue-level vectors by pooling each row, yielding one pooled pairwise-property value per residue. Statistical comparisons between binding and non-binding residue distributions were then reported for each descriptor.

### 2.7 Feature attribution and interpretability

We quantified feature contributions using a two-part attribution approach. First, within-group channel rankings were computed using DeepLIFT-SHAP[39] from Captum(v0.7), applied separately to global scalar, per-residue local, and pairwise feature groups. Multi-dimensional tensors were flattened for attribution and dynamically reshaped within a model wrapper, while non-attributed inputs were held fixed at their input values. For each query protein, 20 baseline proteins were sampled from the test set to reflect the empirical distribution of input features. Per-channel importances were obtained by averaging absolute attributions over valid (annotated) residue positions. Second, to enable cross-group comparison, we computed exact Shapley values for the three feature groups treated as players in a cooperative game. This was done by evaluating the model’s prediction under all 2^3^ = 8 feature coalitions, with missing groups filled by randomly sampled reference proteins. The resulting group-level Shapley values decompose the total prediction contribution additively and provide a common scale for comparison. These were then used to rescale per-channel DeepLIFT-SHAP scores into a unified, cross-group comparable importance measure. All attributions were performed on the corresponding held-out test set for each trained model (LIP or MoRF × AF-CALVADOS, IDPFold2, or STARLING). Robustness across ensemble generators was assessed using Spearman rank correlations of the final feature-importance rankings.

### 2.8 Geometric quality of generated ensembles

Across the tested sources, backbone angle geometry was consistently preserved, but full all-atom viability was rare. IDPFold2 produced the most favorable reconstructed geometries, with 98.04% of covalent bonds within limits, 0.94% clash-free frames, and 0.22% frames passing all geometric checks. AF-CALVADOS showed similar bond accuracy after reconstruction (97.57%) but more frequent steric and Ramachandran violations, whereas STARLING reconstructions showed the largest deviations, particularly for peptide planarity and Ramachandran statistics. We therefore interpret these ensembles primarily through robust ensemble-level descriptors rather than through individual-frame all-atom stereochemical perfection.

## 3 Results

### 3.1 Binding residues occupy distinct ensemble-derived property regimes

Before model training, EnsembleMDP descriptors showed visible separation between annotated binding and non-binding residues across the six dataset-ensemble-source combinations (Methods, and Supplementary Figures S3-S5). This separation was strongest and most consistent for contact-map frequency, which showed the largest Cohen’s *d* and significant discrimination between binding and non-binding residues for both LIP and MoRF annotations across all three conformation-generation methods. For contact-map frequency, Cohen’s *d* was nearly always slightly greater than 1, ranging from 1.06 to 1.32, except for the MoRF STARLING comparison, where it was 0.83.

The next most informative descriptors were dynamic cross-correlation maps (DCCM), solvent-accessible surface area (SASA), backbone *ϕ* entropy, and distance fluctuation. For IDPFold2 and AF-CALVADOS ensembles, DCCM and SASA were consistently the second- and third-ranked properties by effect size after contact frequency. In STARLING ensembles, DCCM and SASA remained highly informative but were often slightly lower in the ranking, commonly around the fourth-largest Cohen’s *d*. Backbone *ϕ* entropy and distance fluctuation followed these leading descriptors for IDPFold2 and AF-CALVADOS, but they were especially important for STARLING, where they frequently provided the second-largest effect sizes. These patterns indicate that binding residues are distinguished not by a single local property but by a reproducible combination of contact propensity, correlated motion, solvent exposure, backbone conformational heterogeneity, and distance-fluctuation behavior, with the precise hierarchy depending on the ensemble source.

### 3.2 Ensemble-derived descriptors improve residue-level prediction

Having established that binding residues occupy distinct ensemble-property regimes, we next asked whether these signals improve supervised residue-level prediction. On proteins shorter than 1024 residues, BindCORE was compared with the corresponding sequence-based baselines on LIP and MoRF binding-site prediction using the metrics defined in Methods.

On the LIP benchmark, BindCORE improved ranking performance over CLIP using ensemble-derived inputs (Table 2; Figure 2). The strongest overall PR-AUC was obtained with IDPFold2 ensembles (AP = 0.4174), compared with 0.3749 for CLIP. IDPFold2 also improved AUC-ROC from 0.7789 to 0.7993 and MCC from 0.3719 to 0.4245. AF-CALVADOS produced a different operating profile, but did not improve over CLIP on the principal evaluation metrics. The ROC curves show that BindCORE variants generally improve residue ranking across a broad range of false-positive rates, whereas the precision-recall curves make the practical gain under class imbalance more apparent, particularly for IDPFold2 on LIP prediction.

**Figure 2.**
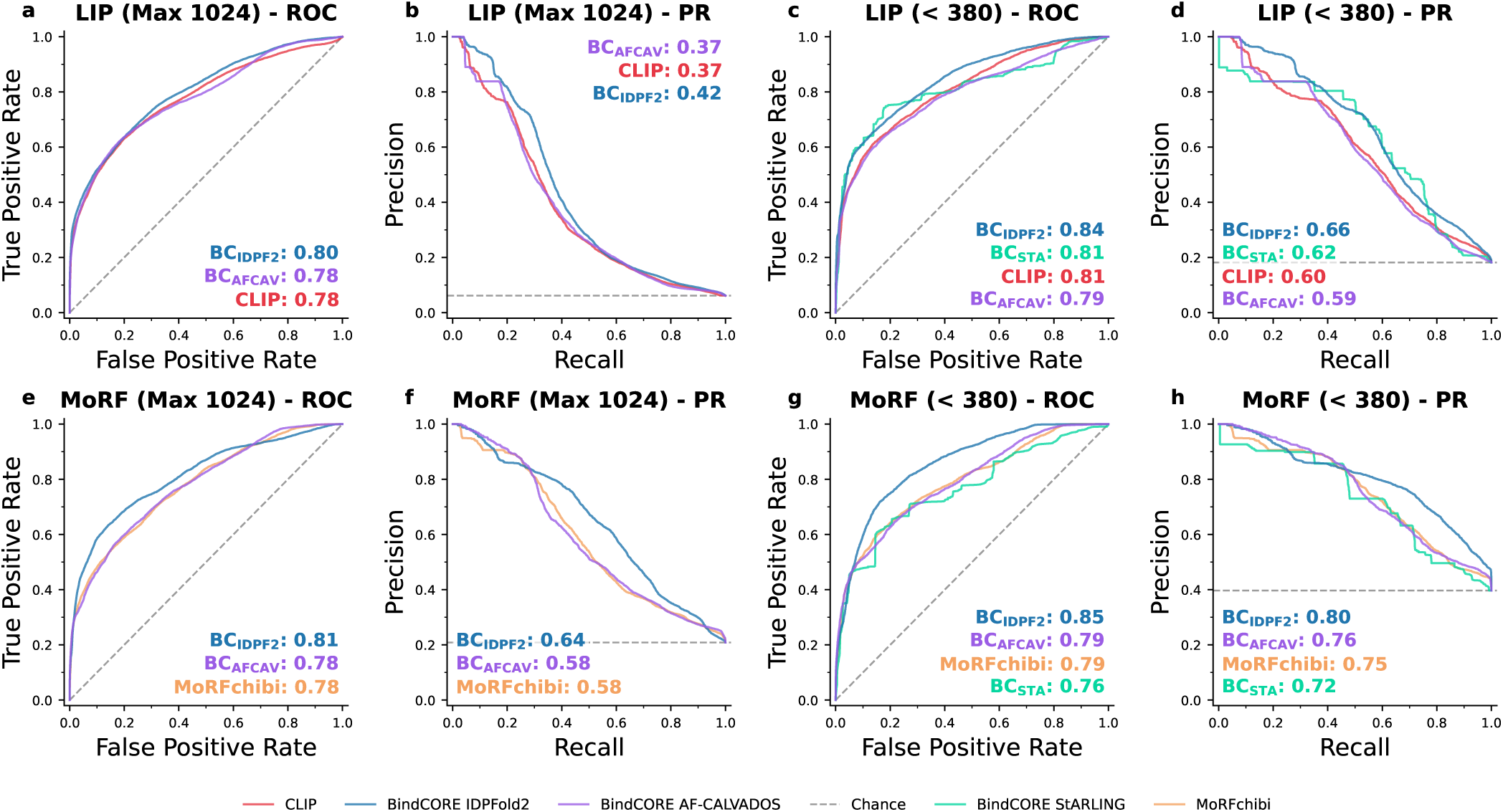
Residue-level performance evaluation across two disorder-centric benchmarks. Each row corresponds to one benchmark dataset: LIP (top) or MoRF (bottom). Each column presents receiver operating characteristic (ROC) curves (left) and precision-recall (PR) curves (right), under two filtering conditions: maximum protein length ≤ 1024 residues (a, b, e, f) and a further restriction to ≤ 380 residues for compatibility with structure-based models such as STARLING (c, d, g, h). Solid colored lines indicate different predictors, including CLIP, Morfchibi 2.0, and BindCORE variants abbreviated as BC-STARLING, BC-IDPFold2, and BC-AF-CALV, while dashed gray lines show random/chance baselines. Boxed annotations display AUC (for ROC) or AP (for PR) values and are positioned to minimize overlap with the curve data.

**Table 1.** Summary of extracted structural and biophysical descriptors and their minimal structural resolution requirements.

| Feature | Granularity | Stored observables | Minimal input |
| --- | --- | --- | --- |
| End-to-end distance ($R_{ee}$) | Global | Mean and std of $R_{ee}^2$ | $C_\alpha$ coordinate |
| Maximum intrachain diameter | Global | Mean, std | $C_\alpha$ coordinate |
| Radius of gyration ($R_g$) | Global | Mean, std | $C_\alpha$ coordinate |
| Gyration eigenvalues and ratios | Global | Mean and std of $\lambda_1, \lambda_2, \lambda_3, \lambda_1/\lambda_2, \lambda_1/\lambda_3$, and $\lambda_2/\lambda_3$ | $C_\alpha$ coordinate |
| Shape anisotropy descriptors | Global | Mean and std of asphericity, prolateness, normalized acylindricity, and relative shape anisotropy | $C_\alpha$ coordinate |
| Flory scaling exponent ($\nu$) | Global | Single fitted scalar | $C_\alpha$ coordinate |
| Dihedral descriptors | Residue-wise ($L$) | $\phi/\psi$ circular mean, std, and entropy ($6L$ values) | Backbone only |
| DSSP-8 secondary-structure propensity | Residue-wise ($L$) | Per-residue DSSP-8 probabilities ($8L$ values) | Backbone only |
| SASA | Residue-wise ($L$) | Absolute and relative mean and std per residue ($4L$ values) | Full-atom |
| Dynamic cross-correlation matrix | Pairwise ($L \times L$) | DCCM matrix | $C_\alpha$ coordinate |
| Distance fluctuation ($CP_{i,j}$) | Pairwise ($L \times L$) | Distance fluctuation matrix | $C_\alpha$ coordinate |
| Contact map frequency | Pairwise ($L \times L$) | Contact probability matrix | $C_\alpha$ coordinate |

**Table 2.** Residue-level performance on proteins shorter than 1024 residues. AP denotes average precision, equivalent to PR-AUC.

| Dataset | Model | MCC | F1 | Precision | Recall | AUC-ROC | AP |
| --- | --- | --- | --- | --- | --- | --- | --- |
| LIP | CLIP | 0.3719 | 0.3776 | 0.5568 | 0.2857 | 0.7789 | 0.3749 |
| LIP | BindCORE AF-CALVADOS | 0.3594 | 0.3681 | 0.5341 | 0.2808 | 0.7797 | 0.3737 |
| LIP | BindCORE IDPFold2 | **0.4245** | **0.4175** | **0.6531** | **0.3069** | **0.7993** | **0.4174** |
| MoRF | Morfchibi 2.0 | 0.4139 | **0.5145** | 0.5969 | **0.4521** | 0.7783 | 0.5823 |
| MoRF | BindCORE AF-CALVADOS | 0.4194 | 0.4429 | 0.7713 | 0.3107 | 0.7797 | 0.5830 |
| MoRF | BindCORE IDPFold2 | **0.4785** | 0.5114 | **0.7961** | 0.3767 | **0.8131** | **0.6403** |

On the MoRF benchmark, BindCORE again improved over the sequence-based reference model (Table 2; Figure 2). The best MCC, precision, AUC-ROC, and AP were achieved with IDPFold2 ensembles (0.4785, 0.7961, 0.8131, and 0.6403, respectively), whereas Morfchibi 2.0 retained the highest F1 score (0.5145) and recall (0.4521). AF-CALVADOS also slightly improved MCC and AP relative to the baseline, but at the cost of substantially lower recall and F1 score. The corresponding curves confirm that the ensemble-based models shift the ranking above the Morfchibi 2.0 baseline for much of the ROC and precision-recall space, while also highlighting that similar AP values can arise from different precision-recall profiles.

### 3.3 STARLING-compatible proteins reveal source-dependent trade-offs

On the STARLING-compatible subset (Table 3), BindCORE models remained competitive with or superior to the corresponding sequence-based baselines, and IDPFold2 was the most consistently strong ensemble source. On the LIP benchmark, BindCORE-IDPFold2 achieved the best MCC (0.5359), F1 score (0.6130), AUC-ROC (0.8428), and AP (0.6623), improving over CLIP on all reported metrics (Table 3; Figure 2). STARLING also improved over CLIP across all metrics, including MCC (0.5353 versus 0.4614), F1 score (0.6076 versus 0.5551), AUC-ROC (0.8115 versus 0.8086), and AP (0.6172 versus 0.5967). The curve-based comparison shows that IDPFold2 provides the most consistent improvement in ranking on this short-protein subset, while STARLING also delivers substantial gains over CLIP across both threshold-dependent and threshold-independent metrics. AF-CALVADOS remained broadly comparable to CLIP but did not match the performance of the other BindCORE variants.

**Table 3.** Residue-level performance on the subset of proteins shorter than 380 residues, enabling comparison with STARLING-derived ensembles. AP denotes average precision, equivalent to PR-AUC.

| Dataset | Model | MCC | F1 | Precision | Recall | AUC-ROC | AP |
| --- | --- | --- | --- | --- | --- | --- | --- |
| LIP | CLIP | 0.4614 | 0.5551 | 0.5784 | 0.5335 | 0.8086 | 0.5967 |
| LIP | BindCORE STARLING | 0.5353 | 0.6076 | **0.6783** | 0.5503 | 0.8115 | 0.6172 |
| LIP | BindCORE AF-CALVADOS | 0.4551 | 0.5418 | 0.6016 | 0.4929 | 0.7889 | 0.5857 |
| LIP | BindCORE IDPFold2 | **0.5359** | **0.6130** | 0.6588 | **0.5732** | **0.8428** | **0.6623** |
| MoRF | Morfchibi 2.0 | 0.4317 | 0.6609 | 0.6507 | **0.6714** | 0.7899 | 0.7511 |
| MoRF | BindCORE STARLING | 0.4342 | 0.4983 | **0.8981** | 0.3448 | 0.7642 | 0.7211 |
| MoRF | BindCORE AF-CALVADOS | 0.4608 | 0.6161 | 0.7763 | 0.5108 | 0.7917 | 0.7580 |
| MoRF | BindCORE IDPFold2 | **0.5323** | **0.6843** | 0.7949 | 0.6008 | **0.8548** | **0.7999** |

On the MoRF benchmark restricted to short proteins, the relative behavior of the input sources was more heterogeneous (Table 3; Figure 2). BindCORE-IDPFold2 provided the best MCC (0.5323), F1 score (0.6843), AUC-ROC (0.8548), and AP (0.7999), while STARLING achieved the highest precision (0.8981). Morfchibi 2.0 retained the highest recall (0.6714), reflecting a more recall-oriented operating point. AF-CALVADOS achieved intermediate performance and remained competitive with the Morfchibi 2.0 baseline in AP (0.7580 versus 0.7511), whereas STARLING underperformed the other ensemble sources on most metrics despite its very high precision. The ROC and precision-recall curves emphasize this heterogeneity: IDPFold2 gives the clearest improvement in threshold-independent ranking, AF-CALVADOS remains competitive across much of the precision-recall space, and STARLING exhibits a markedly more conservative prediction profile. Overall, these results indicate that conformational ensemble descriptors can improve residue-level interaction-site prediction, but the magnitude and nature of the improvement depend on both the benchmark and the ensemble-generation method.

#### 3.3.1 Performance Declines with Protein Length, with Distinct Patterns per Binding Type

For LIPs, all models achieve AUPRC values of roughly 0.80-0.90 on the shortest proteins, with performance declining as longer proteins are included (Figure 3A). IDPFold2 maintains the strongest trajectory across most length thresholds and remains around AUPRC ≈ 0.42 at *k* = 1000 residues, whereas AF-CALVADOS and CLIP converge slightly lower, around AUPRC ≈ 0.38. STARLING can only be evaluated over the shorter-protein range and reaches its evaluation limit at 380 residues. For MoRFs, the performance profile is qualitatively different: all models peak around *k* = 100-200 residues, reaching AUPRC values of 0.90-0.95, before declining as longer proteins are included (Figure 3B). MoRFchibi 2.0 performs strongly on the shortest proteins, consistent with its design as a specialised short-motif predictor, but its performance decreases sharply beyond *k* ≈ 200 residues. By contrast, BindCORE IDPFold2 shows the slowest decline over this longer-protein range and remains around AUPRC ≈ 0.64 at the largest length threshold, while AF-CALVADOS and MoRFchibi 2.0 decline to approximately 0.58.

**Figure 3.**
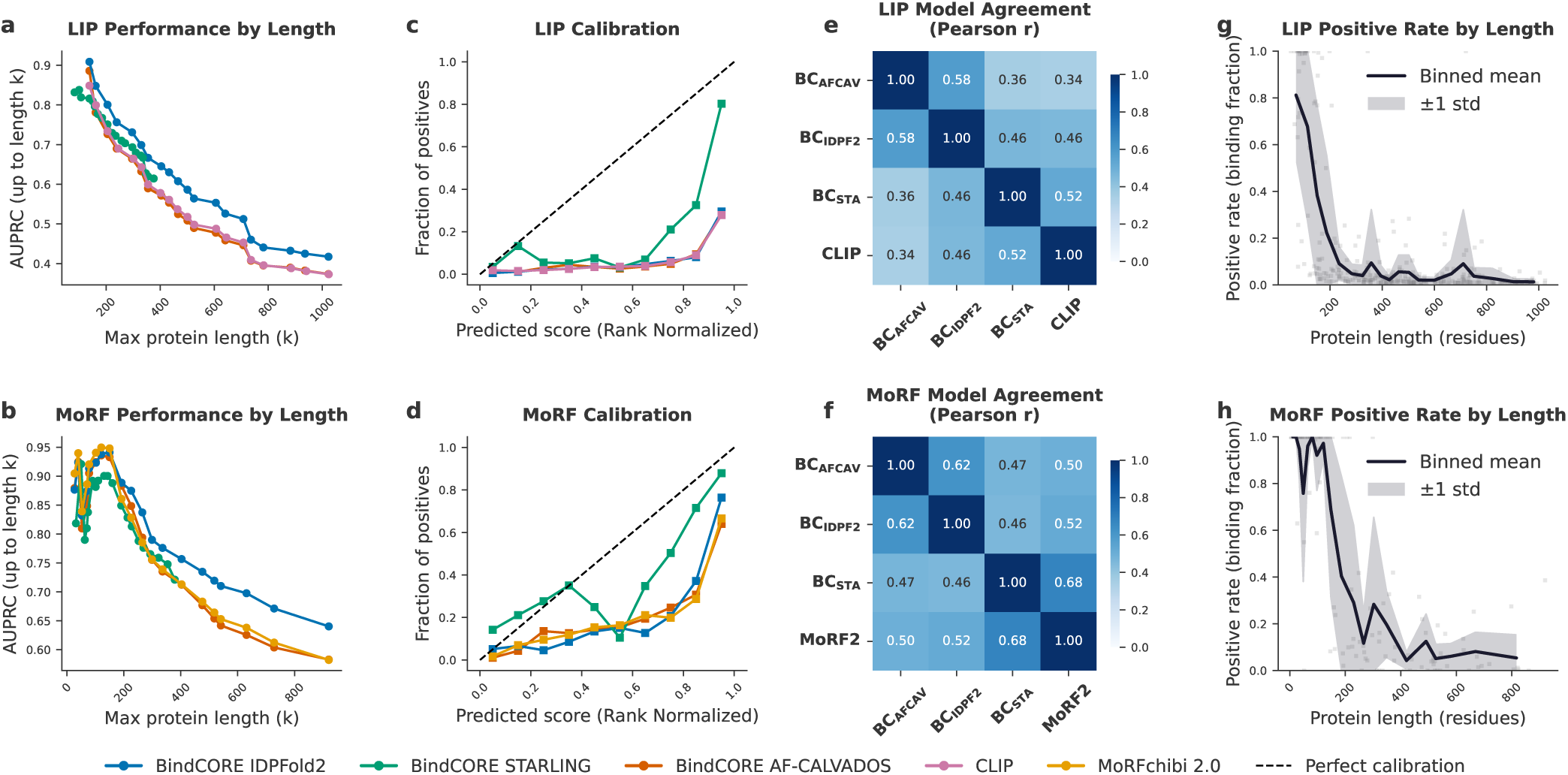
Comparative performance and calibration of binding region predictors. **(a, b)** Area Under the Precision-Recall Curve (AUPRC) evaluated cumulatively up to a maximum protein length threshold (*k*) for LIP and MoRF datasets, respectively. STARLING predictions are limited to sequences shorter than 380 residues. **(c, d)** Reliability diagrams assessing the calibration of rank-normalized prediction scores on the LIP and MoRF datasets. The dashed line indicates perfect calibration. **(e, f)** Pearson correlation matrices quantifying the residue-level agreement between rank-normalized predictions from BindCORE variants and the corresponding sequence-based baseline on the subset of commonly predicted proteins.

#### 3.3.2 Binding Residue Density Decreases Sharply with Protein Length

A prerequisite to interpreting model performance is understanding the distribution of the positive class across protein lengths (Figure 3G, H). For LIPs, the positive rate reaches approximately 0.8 for the shortest proteins and drops rapidly to below 0.2 for sequences longer than 400 residues (Figure 3G). The MoRF dataset exhibits an even more extreme pattern: very short proteins are almost entirely composed of MoRF residues (positive rate ≈ 1.0 for proteins *<* 50 aa), with a steep decline stabilising around 0.2-0.3 for proteins longer than 200 residues (Figure 3H). This length-dependent class imbalance, with high variance throughout both datasets, underscores the fundamental difficulty of benchmarking these models uniformly across the full length spectrum.

#### 3.3.3 STARLING Displays Markedly Better Calibration

Reliability diagrams reveal that AF-CALVADOS, IDPFold2, CLIP, and MoRFchibi are all severely miscalibrated on both datasets: the fraction of true positives remains near zero across most of the predicted score range, rising only sharply at the highest score decile (Figure 3C, D). This indicates that these models concentrate discriminative power in a narrow high-score regime but do not produce scores that can be directly interpreted as probabilities. STARLING, by contrast, exhibits substantially better calibration on both LIP and MoRF datasets. On the LIP test set, STARLING’s fraction of positives reaches ≈ 0.8 at the highest predicted score bin (Figure 3C). On MoRFs, STARLING’s reliability curve follows the perfect calibration diagonal more closely than any other model across the full score range (Figure 3D), suggesting that its ensemble-based features encode more reliable uncertainty estimates about residue binding propensity.

### 3.4 Effect of the Number of IDPFold2 Conformations

Reducing the IDPFold2 ensemble from the default 100 conformations to a single conformation substantially degraded BindCORE performance (Supplementary Figure S6; Supplementary Table S2), confirming that predictive information is distributed across the conformational ensemble rather than captured by a single structure. Performance improved steadily as additional conformations were included and largely saturated between 50 and 100 conformations, indicating diminishing returns beyond this range. These results suggest that relatively large ensembles are beneficial for maximizing predictive accuracy while providing a practical compromise between performance and computational cost. Because ensemble generation also affects inference efficiency, we compared the runtime of the three ensemble-generation methods as a function of protein length (Supplementary Figure S7). IDPFold2 exhibited the highest computational cost, particularly for longer sequences, whereas AF-CALVADOS and STARLING remained substantially faster across their supported sequence-length ranges.

#### 3.4.1 Models Capture Partially Overlapping but Largely Independent Signals

Inter-model score correlations were uniformly moderate across both datasets (Figure 3E, F). For LIPs, the highest agreement was observed between AF-CALVADOS and IDPFold2 (*r* = 0.58), followed by STARLING and CLIP (*r* = 0.52), while AF-CALVADOS and CLIP showed the lowest pairwise correlation (*r* = 0.34). For MoRFs, the strongest correlation was found between STARLING and MoRFchibi 2.0 (*r* = 0.68), whereas IDPFold2 and STARLING showed the weakest agreement (*r* = 0.46). The consistently moderate correlations among all model pairs suggest that the predictors capture partially overlapping but non-identical signals, and that ensemble averaging or stacking could further improve prediction accuracy.

### 3.5 Interpretability analysis reveals a global-to-local hierarchy of conformational determinants

DeepLIFT-SHAP[39] attribution analysis revealed that pairwise features consistently dominate the models’ predictions across both binding-site categories, though with marked dependence on which conformational ensemble generator was used to derive them (Figure 4a,b). For LIP, the dynamic cross-correlation map (DCCM) carried by far the highest mean attribution for BC IDPF2 (∼ 0.30), more than twice that of the next most informative pairwise feature for any flavour, whereas BC AFCAV instead relied predominantly on the contact map (∼ 0.15). A similar but inverted pattern emerged for MoRF, where DCCM became the dominant pairwise feature for BC AFCAV (∼ 0.49), substantially outweighing its importance for BC IDPF2 (∼ 0.16) and BC STA (∼ 0.07). Among global scalar descriptors, shape and size metrics derived from the gyration tensor (Rg *λ*_1_, Rg *λ*_3_, Rg *λ*_1_*/λ*_3_) and end-to-end distance statistics were consistently among the topranked features, but their relative weighting again diverged by ensemble flavour: BC STA assigned markedly higher importance to these descriptors for LIP prediction, while for MoRF, BC AFCAV and BC IDPF2 attributed more weight to maximum end-to-end distance and acylindricity. At the per-residue level, solvent accessibility metrics (SASA, both absolute and relative) and backbone dihedral entropy emerged as the most informative local features across both tasks, with BC IDPF2 showing particularly strong reliance on SASA-related descriptors for MoRF (SASA rel. *µ* ∼ 0.175). Quantifying agreement in feature importance rankings across model flavours (Figure 4c) showed generally weak and, in some cases, negative Spearman correlations, most notably between BC_AFCAV and BC_STA for LIP (*ρ* = −0.62) and between BC_IDPF2 and BC_STA for MoRF (*ρ* = −0.16). This lack of cross-flavour consistency indicates that models trained on ensembles from different conformational generators learn substantially different feature-importance hierarchies (Figure 4), suggesting that the choice of ensemble-generation method shapes not only predictive performance but also the underlying biophysical rationale the model exploits to identify LIP and MoRF binding residues.

**Figure 4.**
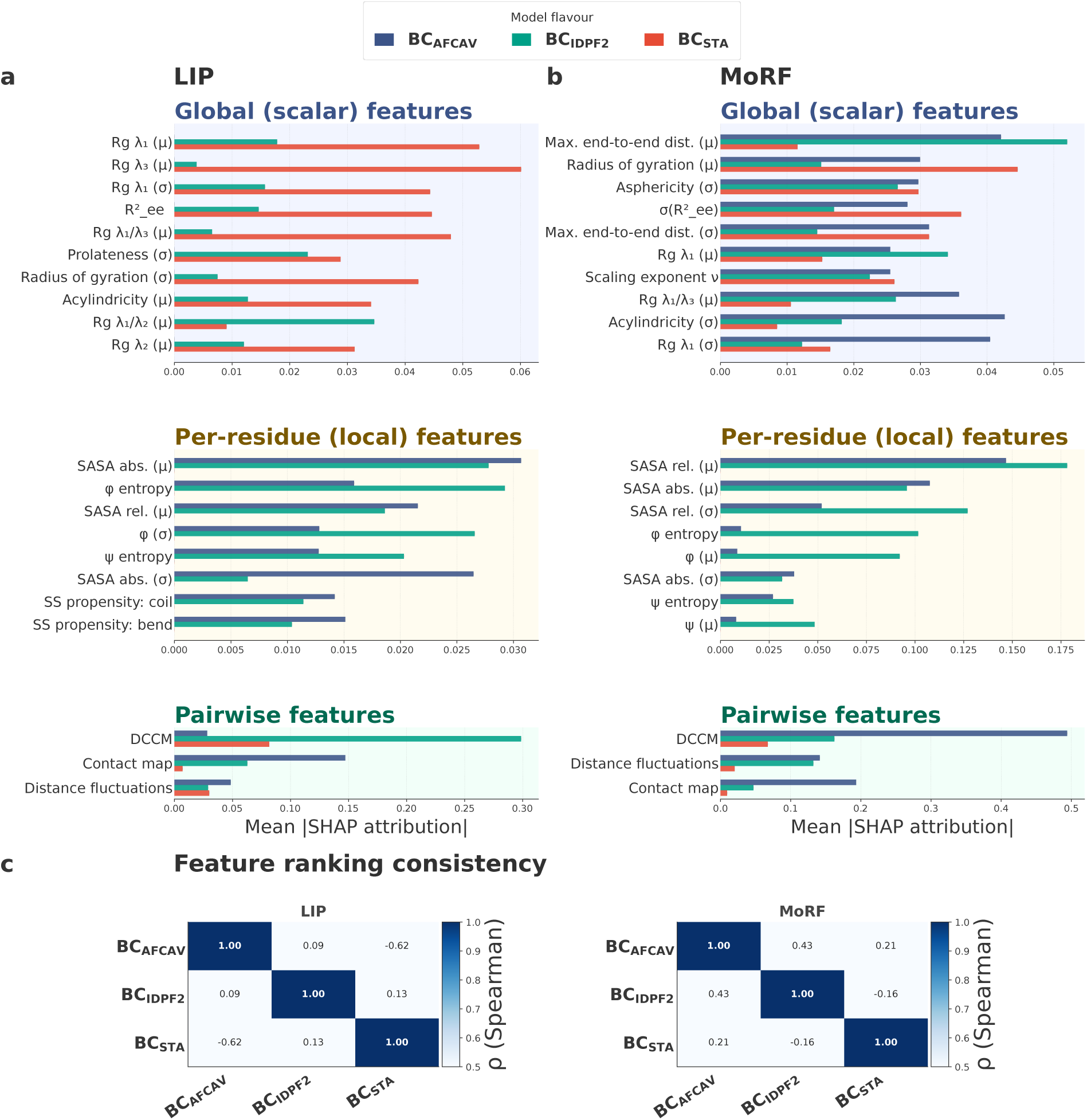
Feature importance analysis of BindCORE. **(a, b)** Top DeepLIFT-SHAP attributions for LIP and MoRF models, grouped as global scalar, per-residue local, and pairwise descriptors and shown separately for AF-CALVADOS, IDPFold2, and STARLING. **(c)** Spearman rank correlations between complete feature-importance rankings across ensemble generators.

## 4 Discussion

This study shows that conformational ensembles add predictive information for residue-level binding-site identification in intrinsically disordered proteins. BindCORE improved ranking performance over CLIP on the LIP benchmark and over MoRFchibi 2.0 on the MoRF benchmark, with the largest gains in average precision under strong class imbalance. These results support the central hypothesis that LIP and MoRF propensity is reflected not only in sequence motifs or evolutionary profiles, but also in the physical organization of the disordered ensemble.

Feature-attribution analysis indicated that pairwise descriptors, especially contact and dynamic cross-correlation maps, contributed strongly to prediction, while global descriptors of size, compaction, and shape and local features such as solvent exposure and backbone torsional statistics provided complementary information. This suggests that binding-competent IDRs occupy ensemble regimes distinct from non-binding disordered regions, with local binding residues embedded in chain contexts that modulate accessibility, pre-organization, intramolecular contacts, and partner engagement.

The ensemble generator also mattered. IDPFold2 gave the strongest and most consistent benchmark results, whereas AF-CALVADOS was generally competitive but often adopted a more conservative operating profile with higher precision and lower recall than the sequence-based baselines. STARLING remains attractive because of its speed and showed strong precision on the short-protein MoRF subset, but sequence-length restrictions, weaker geometric reconstruction, and lower recall limited full-benchmark comparison. BindCORE should therefore be interpreted as an ensemble-aware framework rather than a predictor tied to one sampling method.

Several limitations remain. Reconstructed all-atom frames, especially from coarse-grained or reduced representations, should not be interpreted as individually relaxed structures, although ensemble-level descriptors mitigate this issue. Residue-level annotations are also incomplete and heterogeneous across LIP, MoRF, SLiM, and ProS definitions, which can affect training and evaluation. Finally, differences in feature-importance rankings across ensemble generators may reflect dataset size, feature aggregation, architecture design, or genuine differences in the conformational information encoded by each generator.

Overall, BindCORE establishes a practical route for incorporating conformational ensemble information into IDR interaction-site prediction. As ensemble-generation methods improve, this framework can be extended to larger datasets, additional interaction classes, uncertainty-aware prediction, and mechanistic analyses connecting sequence variation, conformational sampling, and binding function.

## 5 Conflicts of interest

The authors declare that they have no competing interests.

## 6 Funding

This work is supported by the French Agence Nationale de la Recherche (ANR), under grants ANR-22-CPJ2-0075-01 and ANR-24-CE45-4243-01. This work was performed using HPC resources from GENCI–IDRIS (Grant 2026-AD010317569).

## 7 Data and code availability

All code for property computation is available at https://gitlab.inria.fr/delta/ensemblemdp. Code for binding site prediction (LIP and MoRF) and statistical analysis is available at https://gitlab.inria.fr/delta/bindcore.

For direct use, BindCORE is also provided as interactive notebooks (in the colab/ directory of the repository) that predict per-residue LIP and MoRF propensity directly from a protein sequence: a local Jupyter notebook (BindCORE Local.ipynb) and a zero-installation Google Colab notebook (BindCORE Colab.ipynb), the latter runnable at https://colab.research.google.com/github/nbuton/BindCORE/blob/main/colab/BindCORE_Colab.ipynb.

Computed properties for all proteins from IDPFold2, AF-CALVADOS, and STARLING, trained model weights, predictions, interpretability feature importances, and datasets are available in a public Zenodo record at https://zenodo.org/records/20752230.

## 8 Author contributions statement

H.K. conceptualized the core research goals, with N.B. significantly expanding and refining the initial framework. N.B. developed the BindCORE model and, together with L.F.P performed the computational experiments. H.K. supervised the study. All authors contributed to the writing, critical revision, and approval of the final manuscript.

## A Supplementary

### A.1 Supplementary Figures

**Figure S1.**
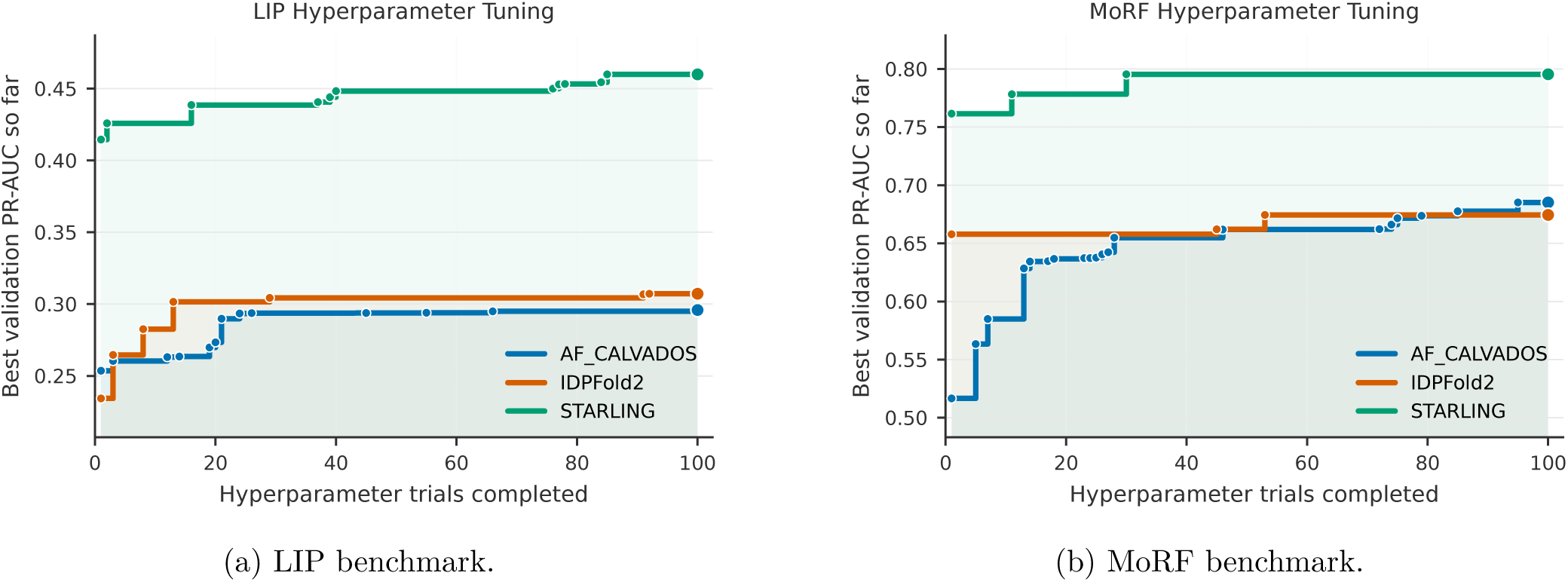
Hyperparameter tuning results for BindCORE on the LIP and MoRF prediction benchmarks. Hyperparameters were optimized with the Tree-structured Parzen Estimator (TPE) sampler implemented in Optuna, using 100 optimization trials for each setting. STARLING-based models are overlaid on the same panels to save space, but STARLING supports sequences only up to 380 residues; therefore, these curves were obtained on the subset of validation proteins with sequence length *<* 380. Consequently, the absolute values of the STARLING curves should not be directly compared with the other curves evaluated on the full validation set; only their overall trend and shape should be interpreted.

**Figure S2.**
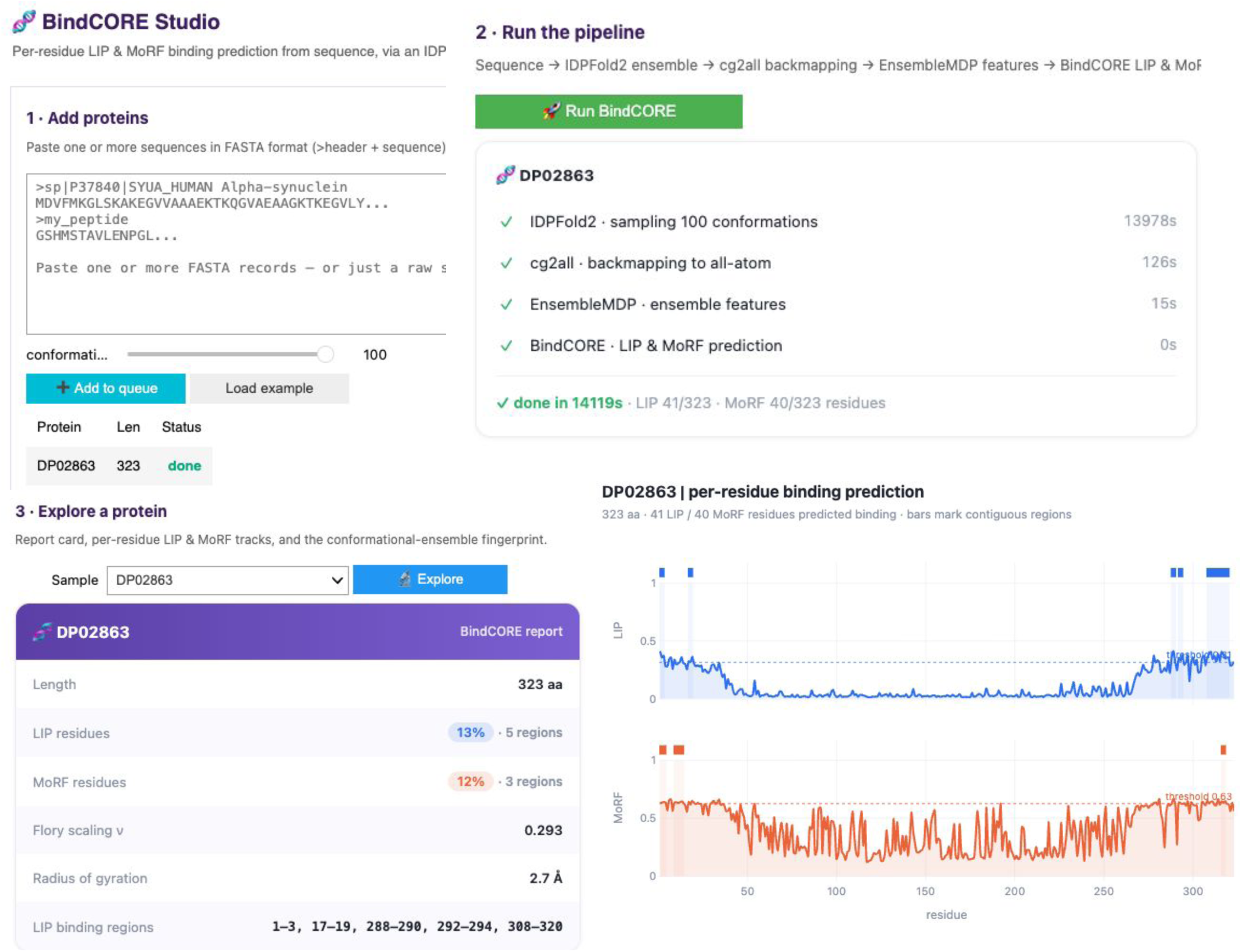
The BindCORE Studio interactive prediction interface. The top panel displays the predicted Linear Motifs (LIPs) and Molecular Recognition Features (MoRFs) profiles across the sequence (e.g., for DP02863), highlighting contiguous binding regions above the threshold. The associated report card displays calculated global biophysical properties, including the Flory scaling exponent (*ν*) and the radius of gyration (*R_g_*), alongside a summary of localized binding regions.

**Figure S3.**
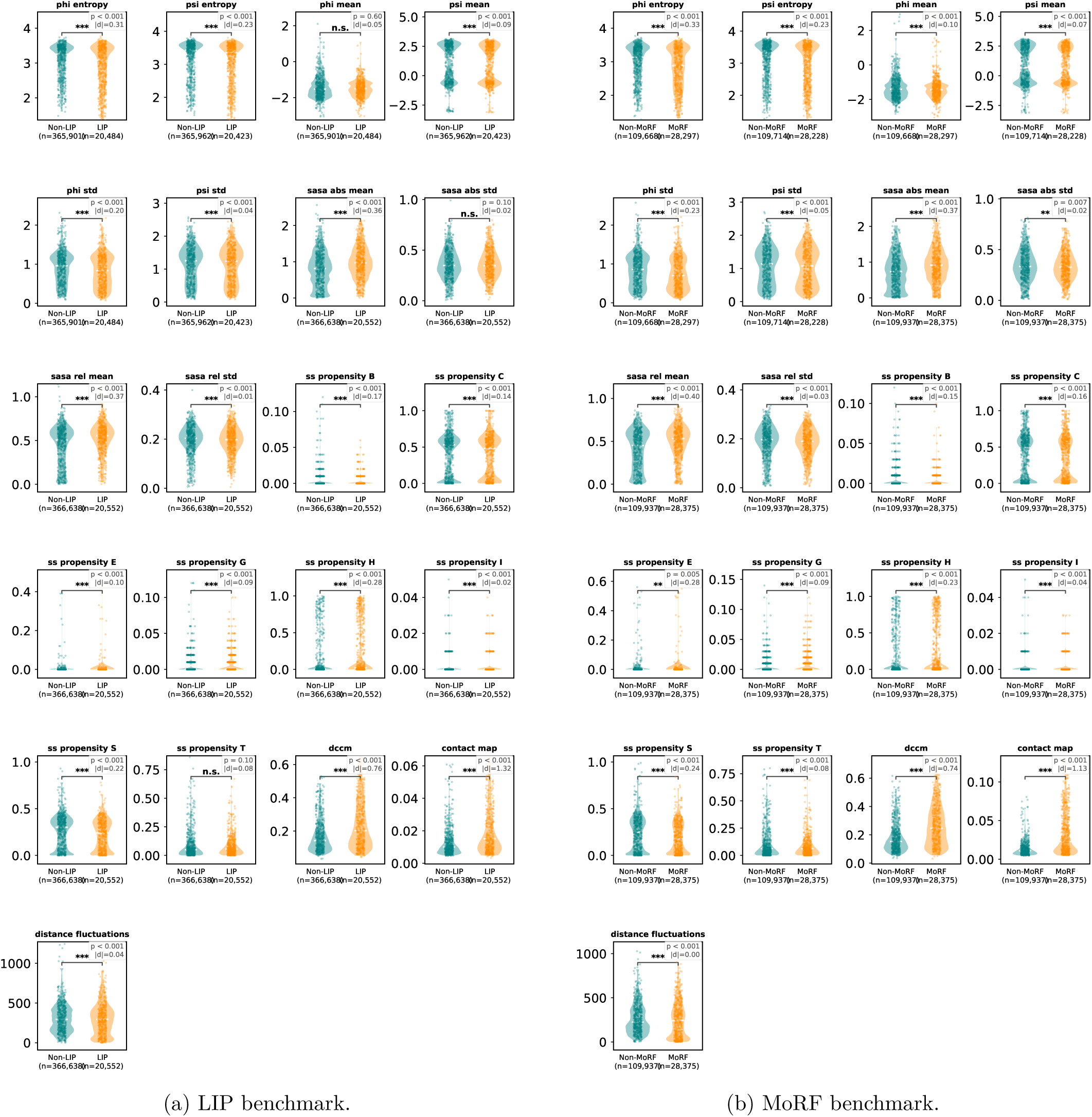
Distribution of ensemble-derived residue properties for annotated binding and non-binding residues using IDPFold2 ensembles. Local descriptors were analyzed directly at residue resolution, whereas pairwise *L L* descriptors were converted to residue-level profiles by pooling each matrix row before plotting. Statistical comparisons between binding and non-binding residues are shown for each property.

**Figure S4.**
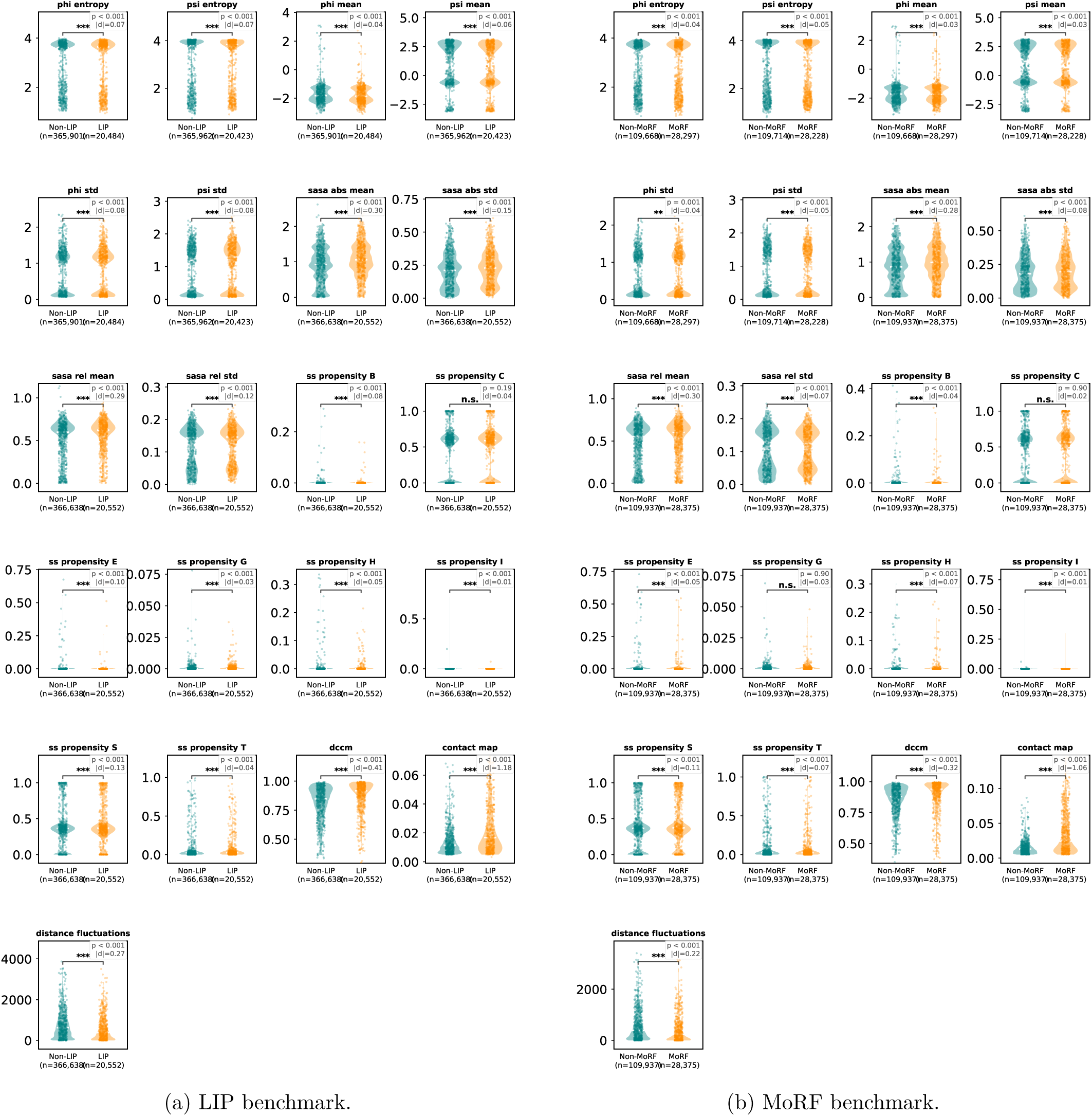
Distribution of ensemble-derived residue properties for annotated binding and non-binding residues using AF-CALVADOS ensembles. Local descriptors were analyzed directly at residue resolution, whereas pairwise *L L* descriptors were converted to residue-level profiles by pooling each matrix row before plotting. Statistical comparisons between binding and non-binding residues are shown for each property.

**Figure S5.**
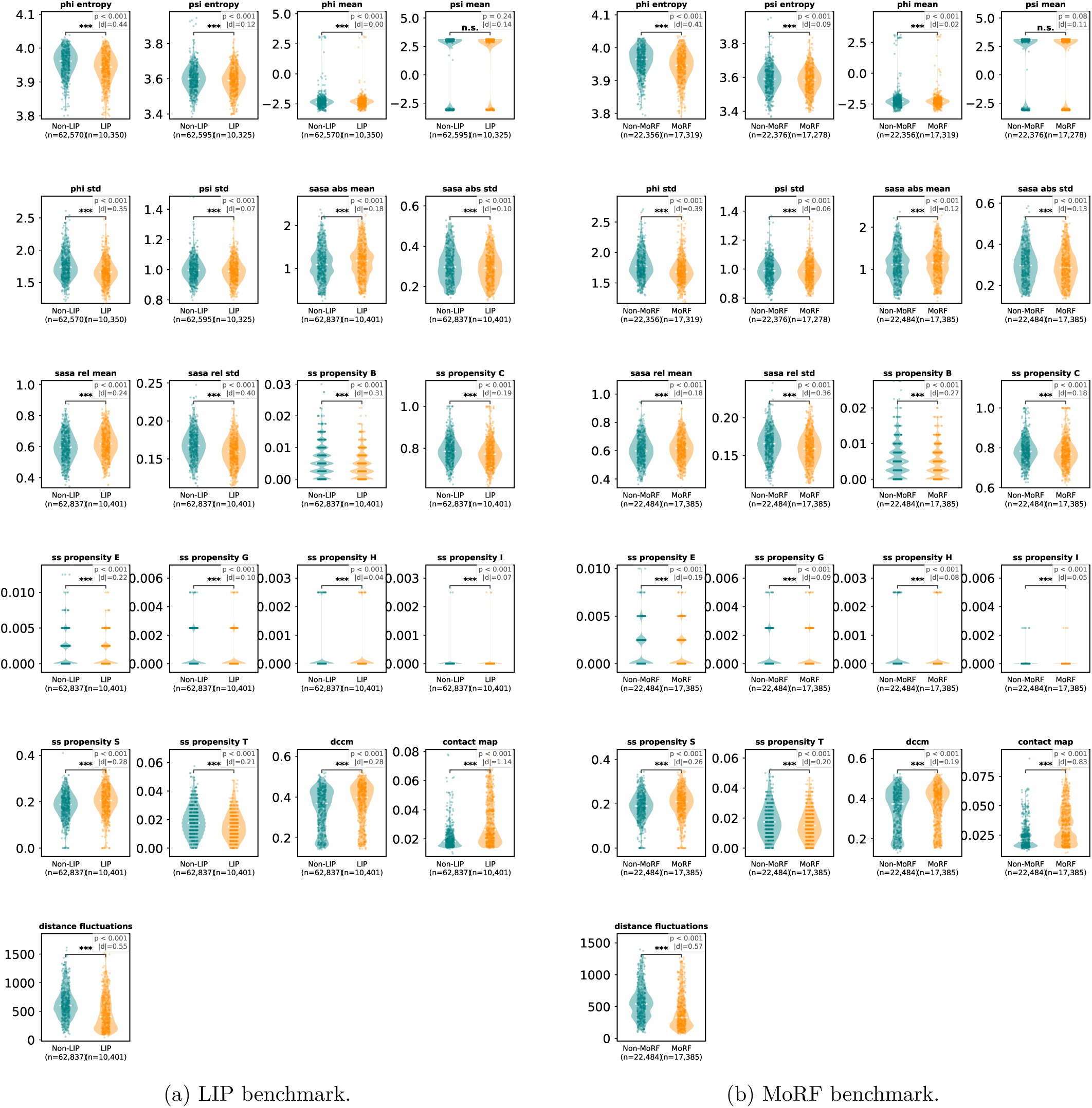
Distribution of ensemble-derived residue properties for annotated binding and non-binding residues using STARLING ensembles. Local descriptors were analyzed directly at residue resolution, whereas pairwise *L L* descriptors were converted to residue-level profiles by pooling each matrix row before plotting. Statistical comparisons between binding and non-binding residues are shown for each property. STARLING panels are restricted to proteins within the method-supported sequence-length range (*L* ≤ 380).

**Figure S6.**
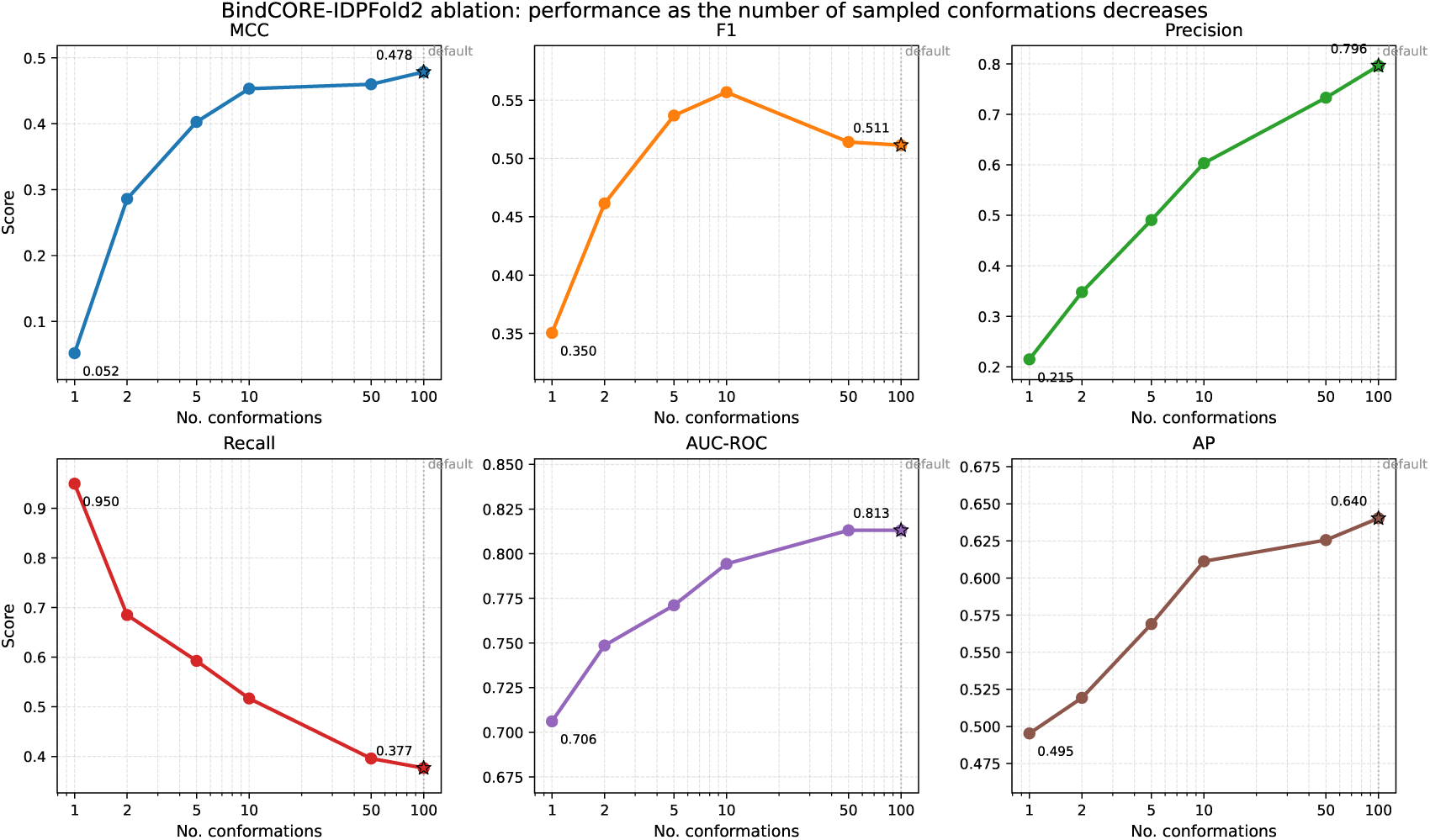
BindCORE-IDPFold2 performance as the number of sampled conformations decreases. Each panel shows one residue-level metric evaluated on the same test set. The default 100-conformation setting is marked at the rightmost point; the single-conformation setting yields a pronounced loss in MCC, AP, and precision, while recall increases substantially.

**Figure S7.**
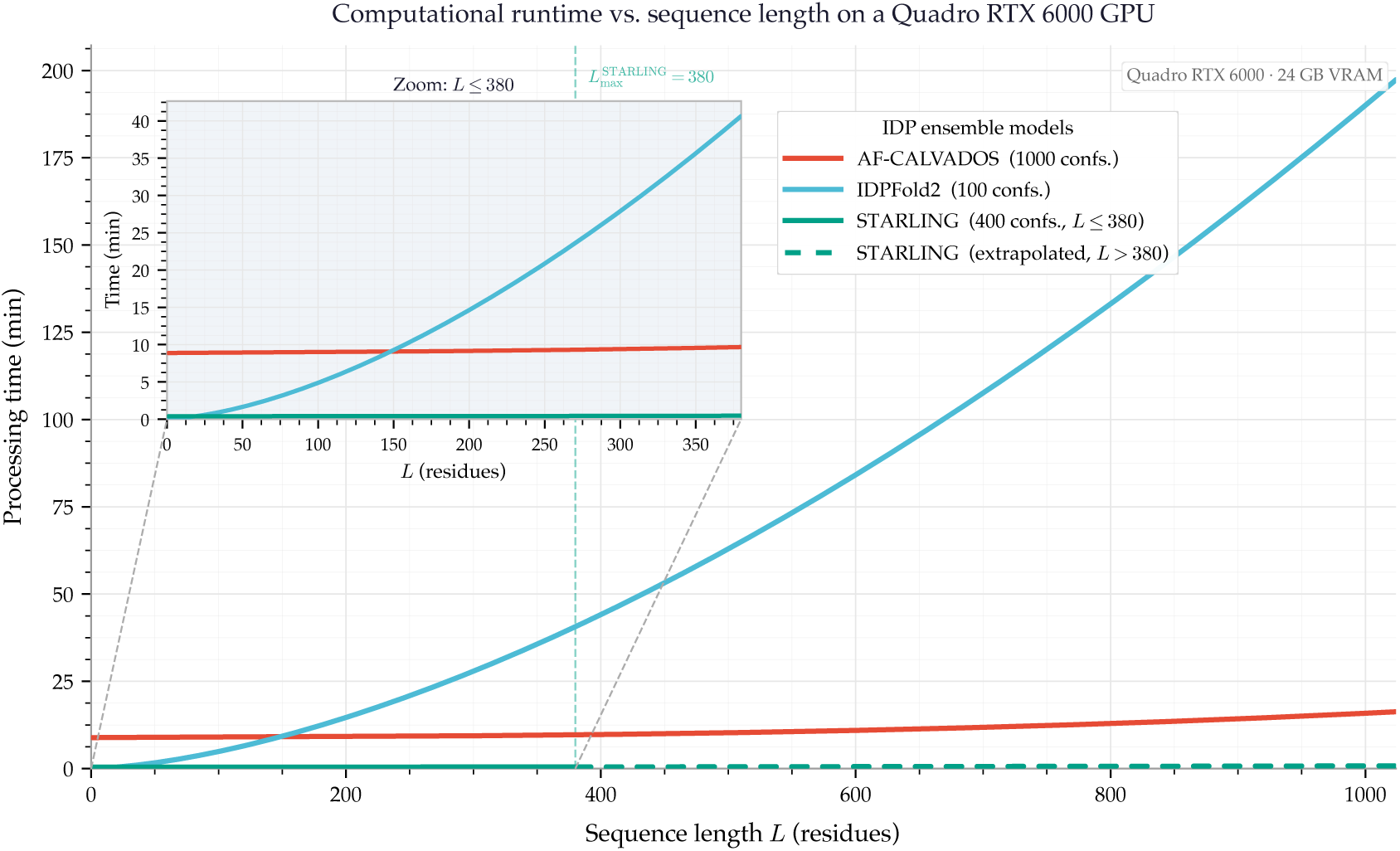
Predicted wall-clock runtime as a function of protein sequence length *L* for three IDP ensemble-generation models benchmarked on an NVIDIA Quadro RTX 6000 GPU (24 GB VRAM). Runtimes are expressed in minutes and were fitted from empirical observations on the target hardware. AF-CALVADOS was run at its default of 1,000 saved conformations; IDPFold2 at 100 conformations; and STARLING at 400 conformations. The solid STARLING curve (teal) covers its supported sequence-length range (*L* 380 residues); the dotted extension beyond *L* = 380 is an extrapolation outside the model’s validated operating range and is shown for reference only. The inset provides a magnified view for short sequences (*L* 380), where IDPFold2 runtimes diverge most clearly from AF-CALVADOS and STARLING.

### A.2 Supplementary Tables

**Table S1.** Statistical summary of the benchmark datasets after sequence-length filtering.

|  | **LIP** | **MoRF** |
| --- | --- | --- |
| Training proteins | 806 | 608 |
| Test proteins | 332 | 125 |
| Median train length | 465 | 375 |
| Median test length | 440 | 359 |
| Positive residues (train) | 5.3% | 11.4% |
| Positive residues (test) | 6.2% | 14.8% |
| Median binding-region length (train) | 7 | 33 |
| Median binding-region length (test) | 8 | 31 |
| Proteins with $\geq 1$ binding region (train) | 100% | 79% |
| Proteins with $\geq 1$ binding region (test) | 100% | 90% |
| Unlabeled residues (train) | 0% | 44.5% |
| Unlabeled residues (test) | 0% | 29.1% |
| Train-test homology filter | 25% (CD-HIT) | 30% + HAM |

**Table S2.** Residue-level performance of BindCORE-IDPFold2 as a function of the number of sampled conformations. The default setting uses 100 conformations. Brier score is omitted.

| conformations | MCC | F1 | Precision | Recall | AUC-ROC | AP |
| --- | --- | --- | --- | --- | --- | --- |
| 1 | 0.0518 | 0.3504 | 0.2148 | 0.9497 | 0.7061 | 0.4953 |
| 2 | 0.2859 | 0.4614 | 0.3479 | 0.6850 | 0.7486 | 0.5193 |
| 5 | 0.4027 | 0.5368 | 0.4906 | 0.5926 | 0.7711 | 0.5690 |
| 10 | 0.4531 | 0.5568 | 0.6033 | 0.5169 | 0.7943 | 0.6113 |
| 50 | 0.4598 | 0.5141 | 0.7329 | 0.3959 | 0.8131 | 0.6256 |
| 100 | **0.4785** | **0.5114** | **0.7961** | 0.3767 | **0.8131** | **0.6403** |

## Notes

https://zenodo.org/records/20752230

### Competing Interest Statement

The authors have declared no competing interest.

## References

[1] Peter E Wright and H Jane Dyson. Intrinsically unstructured proteins: re-assessing the protein structure-function paradigm. Journal of Molecular Biology, 293(2):321–331, 1999.

[2] A Keith Dunker, J David Lawson, Celeste J Brown, Ryan M Williams, Pedro Romero, Jeong S Oh, Christopher J Oldfield, Andrew M Campen, Catherine M Ratliff, Kerry W Hipps, et al. Intrinsically disordered protein. Journal of Molecular Graphics and Modelling, 19(1):26–59, 2001.

[3] Alex S Holehouse and Birthe B Kragelund. The molecular basis for cellular function of intrinsically disordered protein regions. Nature Reviews Molecular Cell Biology, 25:187–211, 2024.

[4] Peter E Wright and H Jane Dyson. Intrinsically disordered proteins in cellular signalling and regulation. Nature Reviews Molecular Cell Biology, 16(1):18–29, 2015.

[5] Robin van der Lee, Marija Buljan, Benjamin Lang, Robert J Weatheritt, Gary W Daughdrill, A Keith Dunker, Monika Fuxreiter, Julian Gough, Joerg Gsponer, David T Jones, et al. Classification of intrinsically disordered regions and proteins. Chemical Reviews, 114(13):6589–6631, 2014.

[6] H Jane Dyson and Peter E Wright. Intrinsically unstructured proteins and their functions. Nature Reviews Molecular Cell Biology, 6(3):197–208, 2005.

[7] Christopher J Oldfield, Yugong Cheng, Marc S Cortese, Pedro Romero, Vladimir N Uversky, and A Keith Dunker. Coupled folding and binding with *α*-helix-forming molecular recognition elements. Biochemistry, 44(37):12454–12470, 2005.

[8] Amrita Mohan, Christopher J Oldfield, Predrag Radivojac, Vladimir Vacic, Marc S Cortese, A Keith Dunker, and Vladimir N Uversky. Analysis of molecular recognition features (MoRFs). Journal of Molecular Biology, 362(5):1043–1059, 2006.

[9] Vladimir Vacic, Christopher J Oldfield, Amrita Mohan, Predrag Radivojac, Marc S Cortese, Vladimir N Uversky, and A Keith Dunker. Characterization of molecular recognition features, MoRFs, and their binding partners. Journal of Proteome Research, 6(6):2351–2366, 2007.

[10] M Madan Babu. The contribution of intrinsically disordered regions to protein function, cellular complexity, and human disease. Biochemical Society Transactions, 44(5):1185–1200, 2016.

[11] Tanja Mittag, Joseph Marsh, Alexander Grishaev, Stephen Orlicky, Hong Lin, Frank Sicheri, Mike Tyers, and Julie D Forman-Kay. Structure/function implications in a dynamic complex of the intrinsically disordered Sic1 with the Cdc4 subunit of an SCF ubiquitin ligase. Structure, 18(4):494–506, 2010.

[12] Benjamin Schuler, Alessandro Borgia, Madeleine B Borgia, Pétur O Heidarsson, Daniel Nettels, and Andrea Sottini. Binding without folding – the biomolecular function of disordered polyelectrolyte complexes. Current Opinion in Structural Biology, 80:102607, 2023.

[13] Sören Von Bülow, Giulio Tesei, and Kresten Lindorff-Larsen. Machine learning methods to study sequence–ensemble–function relationships in disordered proteins. Current Opinion in Structural Biology, 92:103028, 2025.

[14] Zsuzsanna Dosztányi, Bálint Mészáros, and István Simon. ANCHOR: web server for predicting protein binding regions in disordered proteins. Bioinformatics, 25(20):2745–2746, 2009.

[15] Fatemeh Miri Disfani, Wei-Lun Hsu, Marcin J Mizianty, Christopher J Oldfield, Bin Xue, A Keith Dunker, Vladimir N Uversky, and Lukasz Kurgan. Morfpred, a computational tool for sequence-based prediction and characterization of short disorder-to-order transitioning binding regions in proteins. Bioinformatics, 28(12):i75–i83, 2012.

[16] Zhenling Peng, Zixia Li, Qiaozhen Meng, Bi Zhao, and Lukasz Kurgan. Clip: accurate prediction of disordered linear interacting peptides from protein sequences using co-evolutionary information. Briefings in Bioinformatics, 24(1):bbac502, 2023.

[17] Nawar Malhis and Jörg Gsponer. Predicting molecular recognition features in protein sequences with morfchibi 2.0. bioRxiv, pages 2025–01, 2025.

[18] Alexander Miguel Monzon, Paolo Bonato, Marco Necci, Silvio C.E. Tosatto, and Damiano Piovesan. Flipper: Predicting and characterizing linear interacting peptides in the protein data bank. Journal of Molecular Biology, 433(9):166900, 2021.

[19] Hamidreza Ghafouri, Pavel Kaděrávek, Ana M Melo, Maria Cristina Aspromonte, Pau Bernadó, Juan Cortés, Zsuzsanna Dosztányi, Gábor Erdős, Michael Feig, Giacomo Janson, et al. Toward a unified framework for determining conformational ensembles of disordered proteins. Nature Methods, pages 1–15, 2026.

[20] Robert B Best, Nicolae-Viorel Buchete, and Gerhard Hummer. Are current molecular dynamics force fields too helical? Biophysical journal, 95(1):L07–L09, 2008.

[21] Paul Robustelli, Stefano Piana, and David E Shaw. Developing a molecular dynamics force field for both folded and disordered protein states. Proceedings of the National Academy of Sciences, 115(21):E4758– E4766, 2018.

[22] Vincent Schnapka, Tatiana I Morozova, Samiran Sen, and Massimiliano Bonomi. Atomic resolution ensembles of intrinsically disordered proteins with alphafold. Nature Communications, 2026.

[23] Giulio Tesei, Thea K Schulze, Ramon Crehuet, and Kresten Lindorff-Larsen. Accurate model of liquid– liquid phase behavior of intrinsically disordered proteins from optimization of single-chain properties. Proceedings of the National Academy of Sciences, 118(44):e2111696118, 2021.

[24] Giulio Tesei and Kresten Lindorff-Larsen. Improved predictions of phase behaviour of intrinsically disordered proteins by tuning the interaction range. Open Research Europe, 2:94, 2023.

[25] Jerelle A Joseph, Aleks Reinhardt, Anne Aguirre, Pin Yu Chew, Kieran O Russell, Jorge R Espinosa, Adiran Garaizar, and Rosana Collepardo-Guevara. Physics-driven coarse-grained model for biomolecular phase separation with near-quantitative accuracy. Nature computational science, 1(11):732–743, 2021.

[26] Liguo Wang, Christopher Brasnett, Luís Borges-Araújo, Paulo CT Souza, and Siewert J Marrink. Martini3-idp: improved martini 3 force field for disordered proteins. Nature communications, 16(1):2874, 2025.

[27] Junjie Zhu, Sören von Bülow, Hongyi Liu, Kresten Lindorff-Larsen, and Hai-Feng Chen. Extending conformational ensemble prediction to multidomain proteins and protein complex. bioRxiv, pages 2026–01, 2026.

[28] Oufan Zhang, Zi Hao Liu, Julie D Forman-Kay, and Teresa Head-Gordon. Deep learning of proteins with local and global regions of disorder. arXiv preprint arXiv:2502.11326, 2025.

[29] Borna Novak, Jeffrey M Lotthammer, Ryan J Emenecker, and Alex S Holehouse. Accurate predictions of disordered protein ensembles with starling. Nature, pages 1–11, 2026.

[30] Bowen Jing, Bonnie Berger, and Tommi Jaakkola. Alphafold meets flow matching for generating protein ensembles. arXiv preprint arXiv:2402.04845, 2024.

[31] Raphael JL Townshend, Stephan Eismann, and Ron O Dror. Geometric deep learning of RNA structure. Science, 2021. Originally preprint 2019, published 2021.

[32] Albert H Mao, Scott L Crick, Andreas Vitalis, Caitlin L Chicoine, and Rohit V Pappu. Net charge per residue modulates conformational ensembles of intrinsically disordered proteins. Proceedings of the National Academy of Sciences, 107(18):8183–8188, 2010.

[33] Jeffrey M Lotthammer, Garrett M Ginell, Daniel Griffith, Ryan J Emenecker, and Alex S Holehouse. Direct prediction of intrinsically disordered protein conformational properties from sequence. Nature Methods, 21:465–476, 2024.

[34] Michele Invernizzi, Sandro Bottaro, Julian O. Streit, Bruno Trentini, Niccolò Alberto Elia Venanzi, Danny Reidenbach, Youhan Lee, Christian Dallago, Hassan Sirelkhatim, Bowen Jing, Fabio Airoldi, Kresten Lindorff-Larsen, Carlo Fisicaro, and Kamil Tamiola. Advancing protein ensemble predictions across the order–disorder continuum. bioRxiv, 2025.

[35] Lim Heo and Michael Feig. One particle per residue is sufficient to describe all-atom protein structures. bioRxiv, pages 2023–05, 2023.

[36] Yasaman Karami, Elodie Laine, and Alessandra Carbone. Dissecting protein architecture with communication blocks and communicating segment pairs. BMC bioinformatics, 17(Suppl 2):S13, 2016.

[37] John Jumper, Richard Evans, Alexander Pritzel, Tim Green, Michael Figurnov, Olaf Ronneberger, Kathryn Tunyasuvunakool, Russ Bates, Augustin Žídek, Anna Potapenko, et al. Highly accurate protein structure prediction with alphafold. nature, 596(7873):583–589, 2021.

[38] Ashish Vaswani, Noam Shazeer, Niki Parmar, Jakob Uszkoreit, Llion Jones, Aidan N Gomez, L- ukasz Kaiser, and Illia Polosukhin. Attention is all you need. Advances in neural information processing systems, 30, 2017.

[39] Scott M. Lundberg and Su-In Lee. A unified approach to interpreting model predictions. In Proceedings of the 31st International Conference on Neural Information Processing Systems, NIPS’17, page 4768–4777, Red Hook, NY, USA, 2017. Curran Associates Inc.

